## Supplemental Data for "Outer Membrane Vesicles Hijack TIM-1 for Cellular Uptake"

**Extended Data**

**Supplemental Table 1. OMV-binding proteins.** Twenty-six proteins with a normalized relative luminescence unit (RLU) value of >0.01 in both screening replicates.

| Protein ID | RLU (Rep 1) | RLU (Rep 2) |
|---|---|---|
| WBP1 | 0.157 | 0.170 |
| APLP2 | 0.095 | 0.105 |
| TIM-1 | 0.071 | 0.049 |
| TIM4 | 0.057 | 0.046 |
| LSAMP | 0.041 | 0.060 |
| CD300LG | 0.061 | 0.025 |
| NRP1 | 0.049 | 0.036 |
| FCRL2 | 0.056 | 0.025 |
| CD207 | 0.012 | 0.055 |
| LILRA1 | 0.032 | 0.032 |
| SELL | 0.052 | 0.012 |
| CD300A | 0.043 | 0.019 |
| NRP2 | 0.031 | 0.031 |
| KIAA0319L | 0.011 | 0.044 |
| CDCP1 | 0.029 | 0.021 |
| PLXDC1 | 0.022 | 0.028 |
| TREM2 | 0.028 | 0.015 |
| MMP16 | 0.022 | 0.020 |
| CX3CL1 | 0.018 | 0.021 |
| MXRA5 | 0.020 | 0.019 |
| SIGLEC5 | 0.026 | 0.011 |
| SIGLEC6 | 0.025 | 0.011 |
| FCRL6 | 0.017 | 0.017 |
| KIRREL2 | 0.021 | 0.012 |
| CD300C | 0.013 | 0.017 |
| LOC195977 | 0.017 | 0.011 |

**Supplemental Table 2. GO molecular function enrichment for OMV-binding receptors.** Enriched GO terms among the 26 hits are shown with raw P values and false discovery rate (FDR), calculated against the 1,513 proteins screened.

| GO Molecular Function | raw P value | FDR |
|---|---|---|
| Phosphatidylserine binding | 6.26E-06 | 3.43E-03 |
| Modified amino acid binding | 4.21E-05 | 9.21E-03 |
| Heparin binding | 4.08E-06 | 4.47E-03 |
| Glycosaminoglycan binding | 1.28E-05 | 3.50E-03 |
| Sulfur compound binding | 8.11E-06 | 2.96E-03 |

**Supplemental Table 3. OMV size.** Z-average diameter (d. nm) and polydispersity index (PdI) of OMVs as measured by dynamic light scattering.

| Vesicle | Z-Avg | PdI |
|---|---|---|
| *E. coli* K-12 (untreated) | 92.35 | 0.258 |
| *E. coli* K-12 (PMBN) | 89.92 | 0.246 |
| *E. coli* K-12 (proteinase K) | 93.29 | 0.262 |
| *E. coli* K-12 ($Mg^{2+}$) | 112.6 | 0.447 |
| *E. coli* K-12 (Δ*waaC*) | 124.1 | 0.266 |
| *E. coli* K-12 (+*wbbL*) | 133.39 | 0.196 |
| *E. coli* K-12 (+*mcr-1*) | 90.47 | 0.278 |
| LPS (2mg/mL) | 323.2 | 0.382 |
| LPS (0.5mg/mL) | 309.4 | 0.291 |
| LPS (0.125mg/mL) | 5.306 | 0.309 |

**Supplemental Table 4. Bacterial strains used in the study.**

| Bacterial species/strains | Genotype/Other Information |
|---|---|
| *E. coli* BW25113 | Wildtype *E. coli* |
| *E. coli* BW25113 | *E. coli* + pLMG18-ssDsbA: NanoLuc |
| *E. coli* BW25113 Δ*tolQ* | Δ*tolQ*, Keio collection, OMV overproducer |
| *E. coli* BW25113 Δ*tolQ* | Δ*tolQ* + pACYC117-*wbbL* |
| *E. coli* BW25113 Δ*waaC* | ΔwaaC, Keio collection, truncated LPS |
| *E. coli* BW25113 Δ*tolQ* | Δ*tolQ* + pBAD24-*mcr-1* |
| *F. nucleatum* ATCC 23726 | Wildtype Fn23726 |
| *E. coli* (O157:H7) ATCC 35150 | Wildtype enterohemorrhagic *E. coli* (EHEC) |
| *A. baumannii* ATCC 17978 | Wildtype Ab17978 |
| *P. aeruginosa* PA14 | Wildtype PA14 |

**Supplemental Table 5. Cancer cell lines used for this study.**

| **Cell Line** | **Genotype/Other Information** |
|---|---|
| A549 (ATCC) | Human lung epithelial cell line |
| A549 (ATCC) | A549-Vector Control-pBac |
| A549 (ATCC) | A549-TIM-1-pBac |
| A549 (ATCC) | A549-TIM-4-pBac |
| A549 (ATCC) | A549-APLP2-pBac |
| A549 (ATCC) | A549-NRP1-pBac |
| Caco-2 (ATCC) | Human colon epithelial cell line |
| Caco-2 (ATCC) | Caco-2-Vector Control-pBac |
| Caco-2 (ATCC) | Caco-2-TIM-1-pBac |
| Caco-2 (ATCC) | Caco-2-TIM-4-pBac |
| Caco-2 (ATCC) | Caco-2-APLP2-pBac |
| Caco-2 (ATCC) | Caco-2-NRP1-pBac |
| THP-1 (ATCC) | Human monocyte cell line |
| THP-1 (ATCC) | THP-1-Vector Control-pBac |

| | |
|---|---|
| THP-1 (ATCC) | THP-1-TIM-1-pBac |
| 769-P (ATCC) | Human kidney epithelial cell line |
| ACHN (ATCC) | Human kidney epithelial cell line |
| IGROV-1 (DCTD) | Human ovary epithelial cell line |
| Suit-2 (JCRB) | Human pancreas epithelial cell line |
| Huh-7 (JCRB) | Human liver epithelial cell line |
| SNU-449 (ATCC) | Human liver epithelial cell line |
| 786-O (ATCC) | Human kidney epithelial cell line |
| HCC1534 (UTSW) | Human lung epithelial cell line |
| A-704 (ATCC) | Human kidney epithelial cell line |
| Cal-51 (DSMZ) | Human breast epithelial cell line |
| FU97 (JCRB) | Human stomach epithelial cell line |
| HeLa (ATCC) | Human cervix epithelial cell line |
| HCT-116 (ATCC) | Human colon epithelial cell line |
| HT-29 (ATCC) | Human colon epithelial cell line |

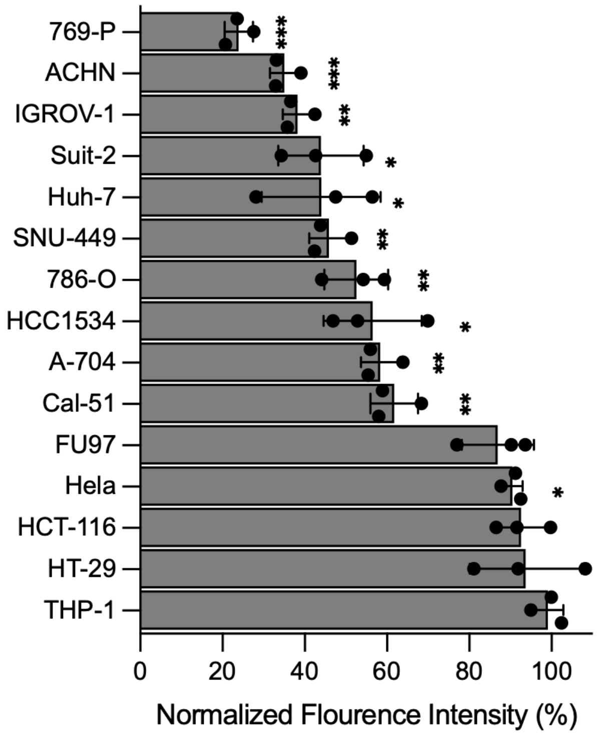


**Supplemental Figure 1. Anti–TIM-1 antibody reduces OMV uptake across TIM-1–expressing cell lines.** Cells were treated with anti–TIM-1 monoclonal antibody (10 µg/mL) or isotype control 30 mins before adding DiO-labeled *E. coli* K-12 OMVs. Uptake was quantified by flow cytometry after 2 hours with trypan blue quenching and normalized to isotype (= 100%). Bars show mean ± std deviation. TIM-1–positive lines (769-P, ACHN, IGROV-1, Suit-2, Huh-7, SNU-449, 786-O, HCC1534, A-704, Cal-51, Caco-2, A549) showed ≥40% reduction with anti–TIM-1, whereas TIM-1–negative lines (HeLa, HCT-116, HT-29, THP-1) were largely unaffected. Bars represent mean ± s.d, n = 3, *** $P < 0.001$, ** $P < 0.01$, * $P < 0.05$ compared to an isotype control, using one-sample t-test vs 100% (two-sided).

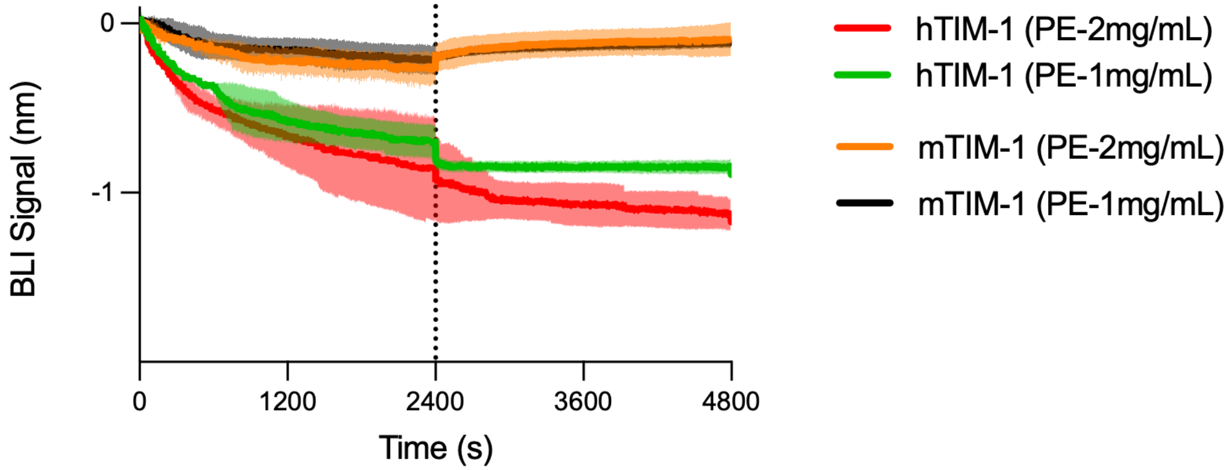


**Supplemental Figure 2. Human TIM-1(hTIM-1) interacts with phosphatidylethanolamine (PE).** PE binding is observed for hTIM-1 but not mouse TIM-1 (mTIM-1) as detected by BLI. Solid line represents the mean and the shaded band is ± s.d. of at least two biological replicates

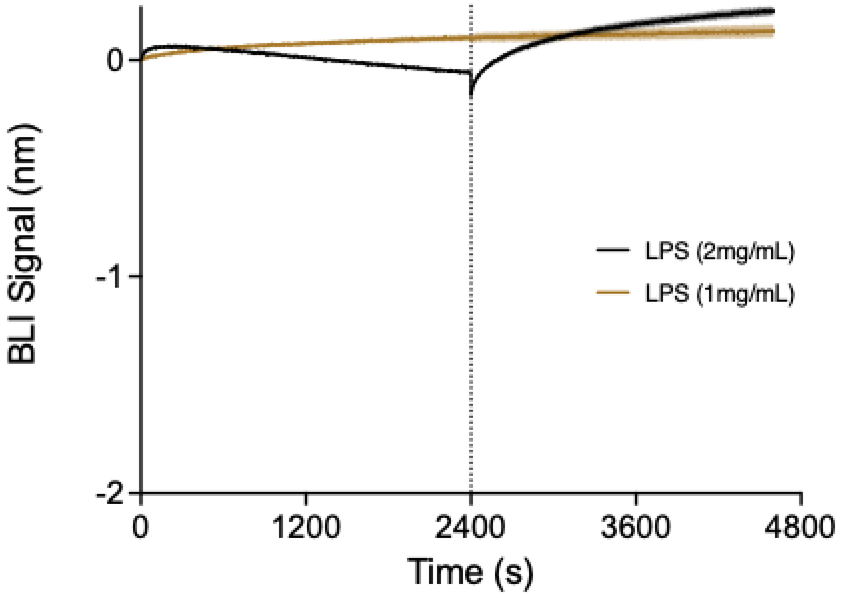

**Supplemental Figure 3. LPS does not indiscriminately bind receptors.** Human EGFR did not demonstrate binding to LPS at concentrations shown to interact with TIM-1. Solid lines represent the mean and the shaded band is ± s.d. of at least two replicates.

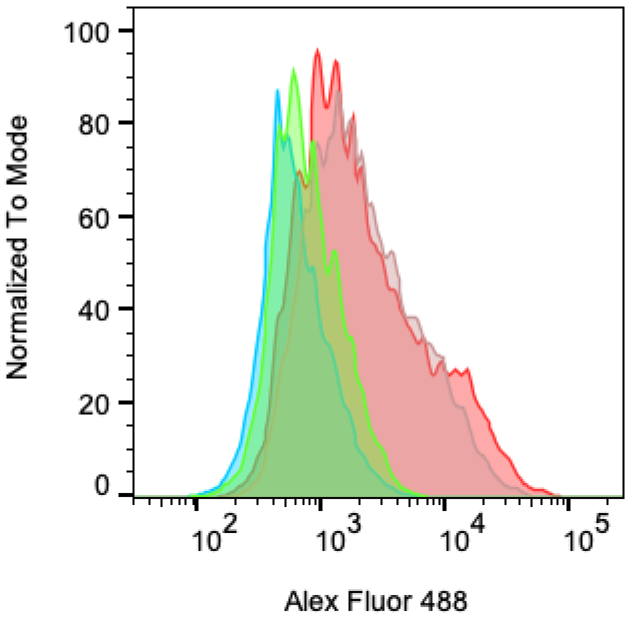


| | Cell Line | Treatment | Vector | Median : Alexa Fluor 488-A |
|---|---|---|---|---|
| | A549 ΔTim-1 | Isotype | Tim-1 (wildtype) | 716 |
| | A549 ΔTim-1 | Anti-Tim-1 | Tim-1 (wildtype) | 1833 |
| | A549 ΔTim-1 | Isotype | Tim-1 (AAAA) | 569 |
| | A549 ΔTim-1 | Anti-Tim-1 | Tim-1 (AAAA) | 1681 |

**Supplemental Figure 4. Surface levels of wildtype and mutant TIM-1.** Overexpression of wildtype TIM-1 and W112A/F113A/N114A/D115A TIM-1 in A549 cells stained with Anti-TIM-1 antibody (P365D) and analyzed with flow cytometry to determine surface levels.

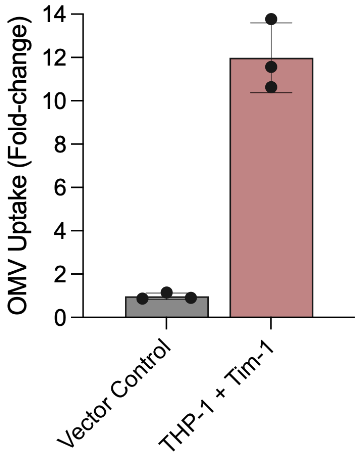


**Supplemental Figure 5. Overexpression of TIM-1 in THP-1 monocytes.** Undifferentiated THP-1 monocytes are treated with 50 μg/mL of DiO stained *E. coli* OMVs for 2 hours and fluorescence (488) determined with flow cytometry. Data is normalized to wildtype THP-1 monocytes.
